# MAGE-3 peptide amphiphile micelle vaccine promote anti-tumor immunity in mice with stomach cancer

**DOI:** 10.1101/609214

**Authors:** Joseph Windberg, Rui Zhang

## Abstract

Nanoparticles as a vaccine carrier can protect antigen from enzymatic hydrolysis, enhance immunogenicity, is a kind of great potential for development of new vaccine carriers. In this study, a nanometer vaccine loaded with CD4+ & CD8+ T cell epitope MAGE-3 polypeptide antigen was prepared to investigate its related properties and anti-tumor immunity. Methods: the use of self-assembly technology to prepare polypeptide / Chit2DC (chitosan - deoxycholate) drug micelles, transmission electron microscopic morphology, fluorescence spectrophotometry to calculate the loading rate, drug loading, and drug release rule. Flow cytometric detection of DC (dendritic cells) on the phagocytic rate of the drug, enzyme-linked immunosorbent spot test (ELISPOT) and cytotoxicity assay MAGE-3 polypeptide nanometer vaccine activation status of the body’s cellular immune response. In vivo tumor suppressor effect was observed in animals. Results: the peptides /Chit2DC micelles were prepared successfully. the drug encapsulation efficiency was about 37% and the drug loading was 17%. Drug-loaded nanoparticles polypeptide at pH 7.14 of the "cancer" ELISPOT and cytotoxicity experiments show that MAGE-3 polypeptide nanometer vaccine can activate the immune response in vivo to produce CTL against MAGE-3, specifically killing tumor cells expressing MAGE-3. In vivo tumor inhibition experiments showed that the relative tumor inhibition rate of polypeptide nanoparticles group was 37.181%.

## Introduction

In recent years, the rapid development of tumor vaccine, polypeptide vaccine because of its strong specificity, low side effects, ease of synthesis and preparation has become one of the hot spots of tumor vaccine research. However, low immunogenicity polypeptide vaccine, susceptible to enzymatic hydrolysis in vivo, low bioavailability, poor biological stability, the need for a suitable carrier material to overcome its shortcomings. Nanomaterials as a polypeptide vaccine Vector has made great progress, which can protect the antigen, improve immunogenicity; and because of the relationship between the particle size can be directly drained to regional lymph nodes, with a certain lymphatic targeting[1–8]. However, some of the traditional nanomaterials are poor biocompatibility, the preparation process is complex, requires some chemical mediators involved in the synthesis, prone to damage to the biologically active molecules. Molecular self-assembly is a new concept proposed in recent years, which is the use of non-covalent intermolecular forces, the construction of drug-loaded micelles, can effectively protect the biologically active molecule from destruction[9–11]. In this experiment, the polypeptide nanoparticles were constructed by the method of molecular self-assembly, and the in vitro immune mechanism and the inhibitory effect on tumor-bearing mice were studied.

## Results

Figure 1 shows the peak between 0.16 ~ 2.15 for deoxycholic acid magnetic resonance peak, peak between 3 to 5 for the chitosan backbone magnetic resonance peak, confirmed deoxycholic acid and chitosan conjugate.

**Figure 1.**
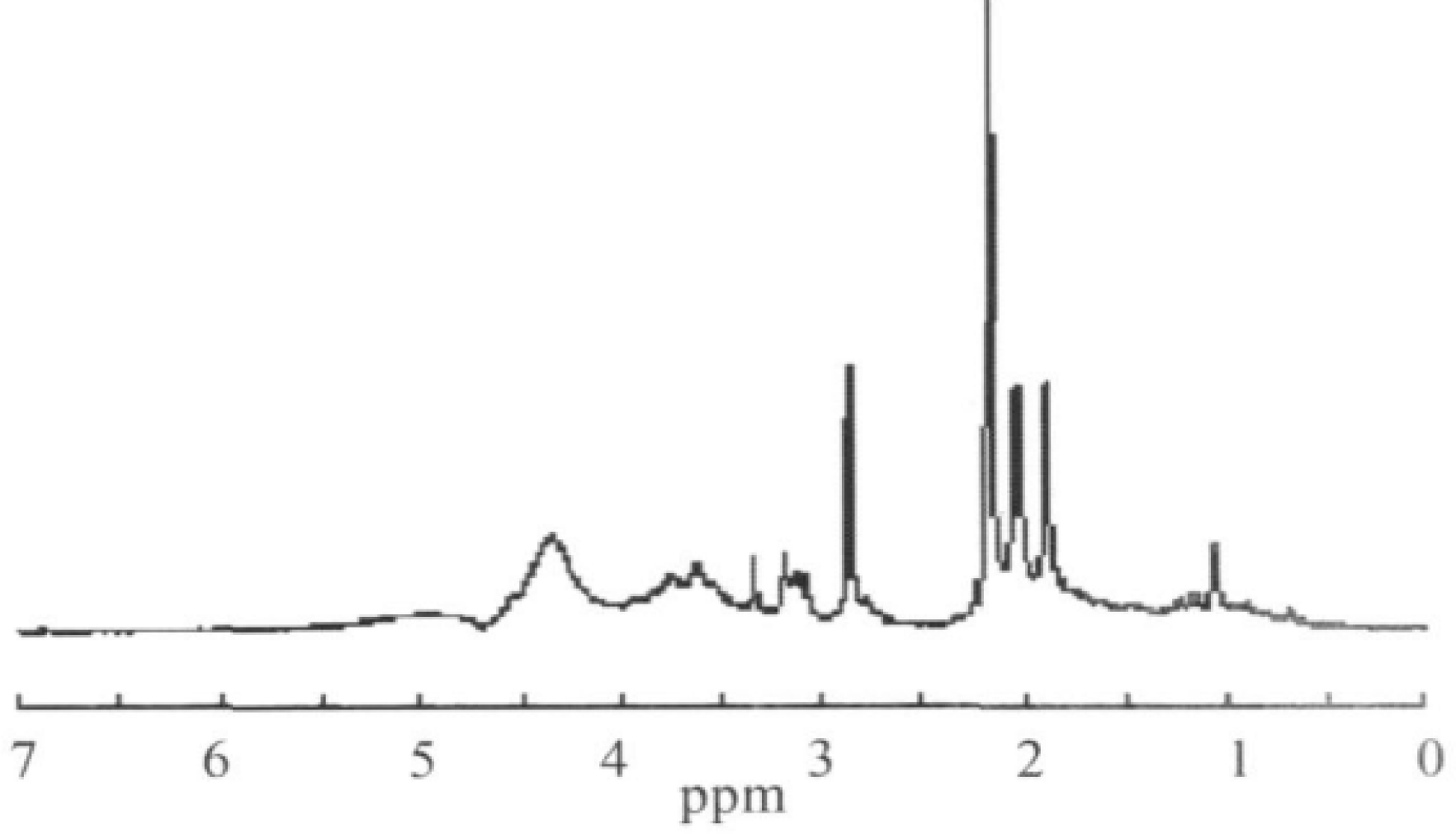
NMR Spectra of peptide amphiphile.

TEM observed Chit2DC micelles were 30 ~ 50 nm particles, round or round, less uniform size, See Figure 2.

**Figure 2.**
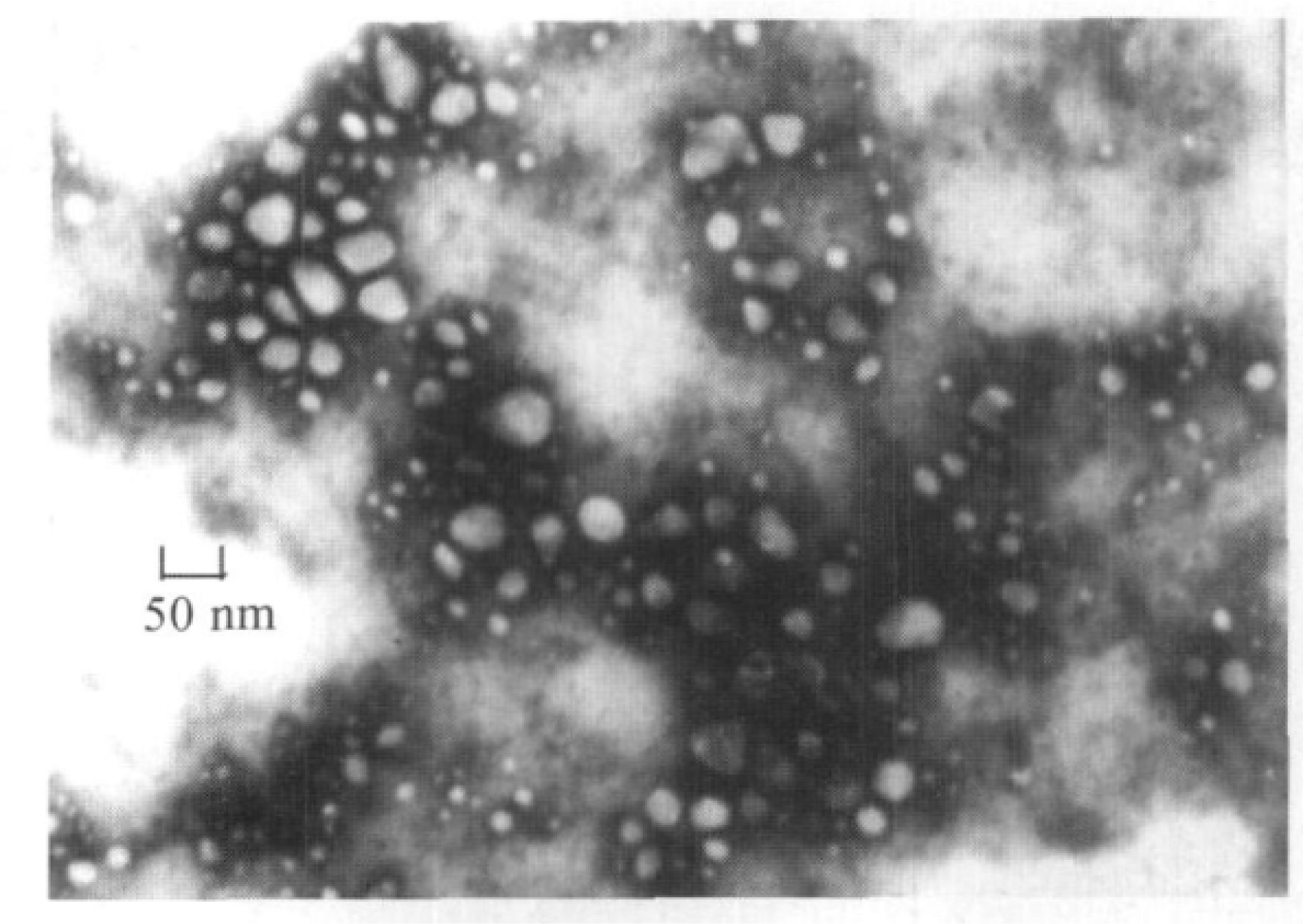
TEM images of MAGE-3 peptide amphiphile micelles

Fluorescence spectrophotometry standard curve, the result is: I = - 3110 + 74120C (R = 0.1998). According to the curve, the load can be measured was 37.13%, the drug loading was 1710%. 214 MAGE-3 polypeptide vaccine in vitro release behavior fluorescence spectrophotometry work curve obtained results: I = - 0186 + 43145C (R = 0.1998).Plotted release profile according to the concentration of the polypeptide required by the standard curve.

Shown in Figure 3A, the drug-loaded nanoparticles slow release of the polypeptide, after about 48 h release completely; in lysozyme solution, shown in Figure 3B, the polypeptide of about 24 h release platform. Lysozyme has a certain role in promoting the release of chitosan nanoparticle drug-loaded micelles, but it still maintains a certain sustained-release effect. The phagocytosis of dendritic cells of the 215 MAGE-3 polypeptide nanometer vaccine collected suspended and semi-adherent cells after fixation electron microscopy of the cell surface with Burr protrusions, showing a typical DC form. By Flow Cytometry found that DC uptake of FITC-MAGE-3 polypeptide nanometer vaccine is higher than the free FITC-MAGE-3 polypeptide vaccine uptake rate (P < 0.105), was dose-dependent (10 µg group: 50.12% vs11.112%; 30 µg group: 62.14% vs12.316%; 50 µg group: 66.18% vs13.410%; 70µg group: 72.14% vs14.014%. Wherein PBS negative control group was 2.14%).

**Figure 3.**
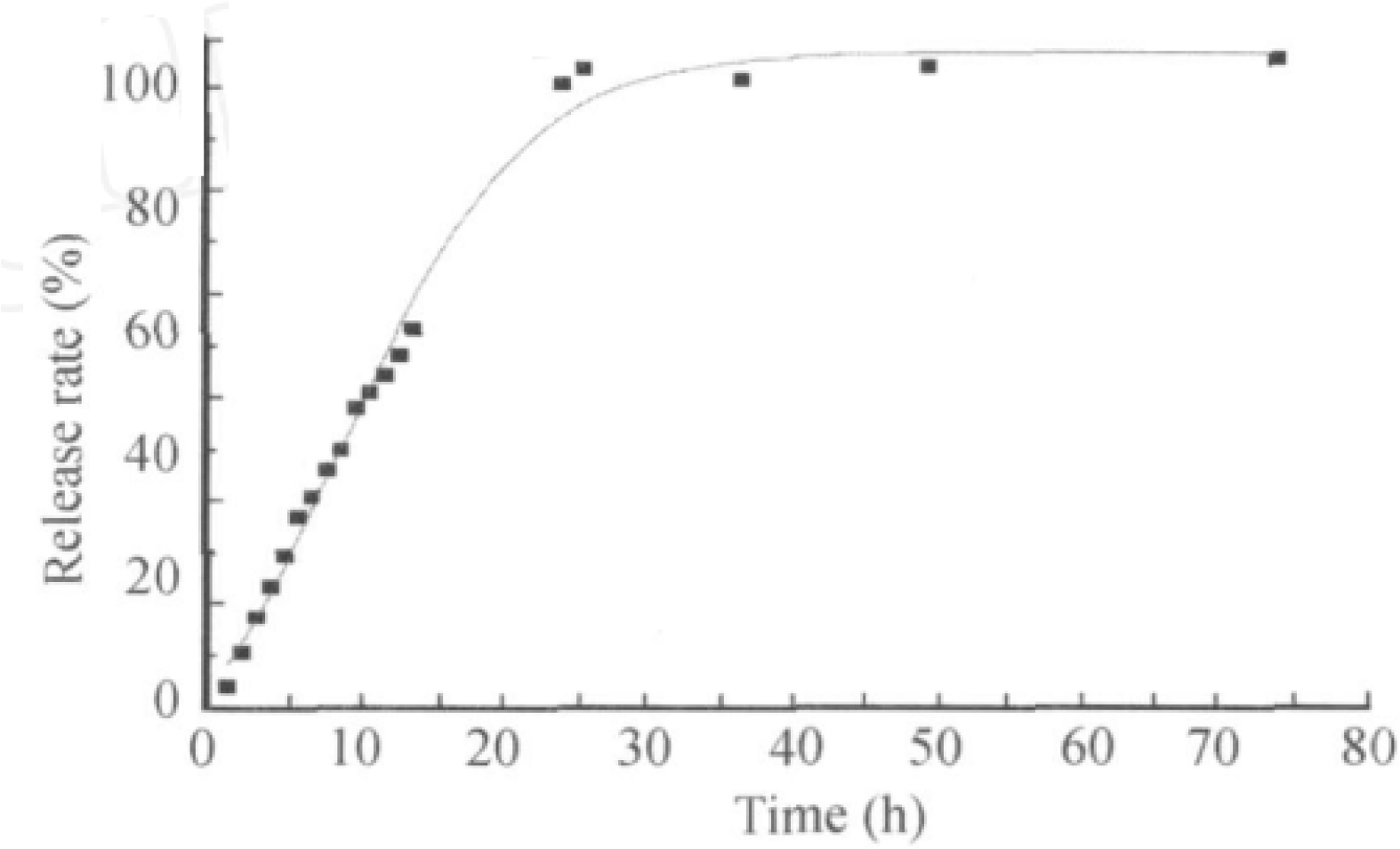
Releasing kinetic study indicates the drug was released within 30 hours.

**Figure 4.**
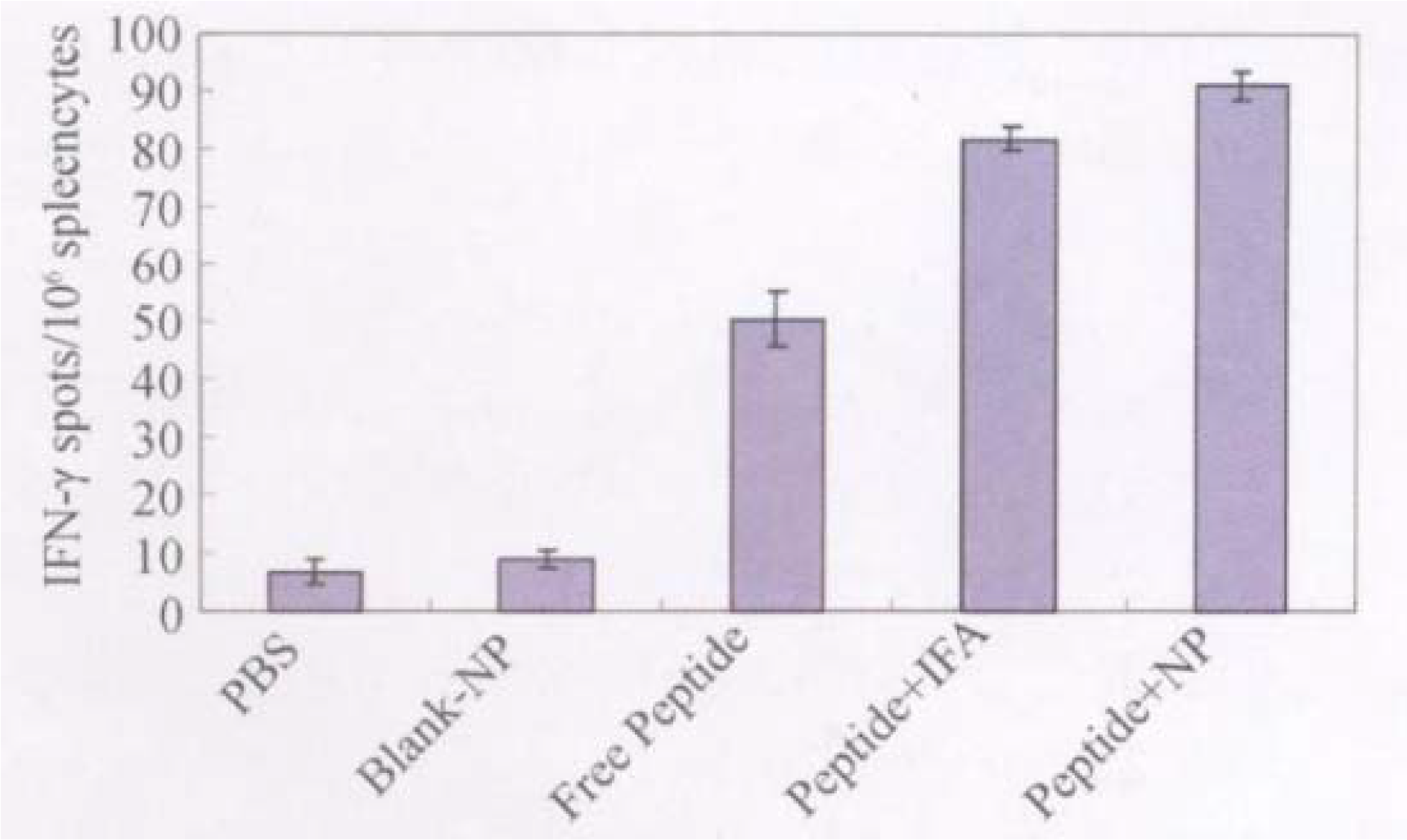
IFN secretion from cells activated with Mage-3 peptide.

The number of specific T-lymphocytes of ifn2î (Ifn2î) secreted by the peptide nanocomposite was higher than that of the control nanocomposite and PBS (P < 0.105), and the number of splenic T-lymphocytes in the peptide nanocomposite was statistically significant compared with that of the polypeptide adjuvant (P = 0.1024).Free MAGE-3 polypeptide, spleen polypeptide + IFA MAGE-3, MAGE-3 polypeptide nanometer vaccine immunized mouse lymphocytes capable of specifically killing the expression of MAGE-3 MFC cells, wherein the polypeptide nanometer different target cell ratio of the vaccine group of mouse spleen lymphocytes specific killing capacity higher than the polypeptide group, the blank nanometer group and PBS group (P < 0.105). The blank nanometer group and PBS group immunized mouse spleen lymphocytes expressing mage-3 positive MFC cells killing effect are weak, shown in Figure 5.

**Figure 5.**
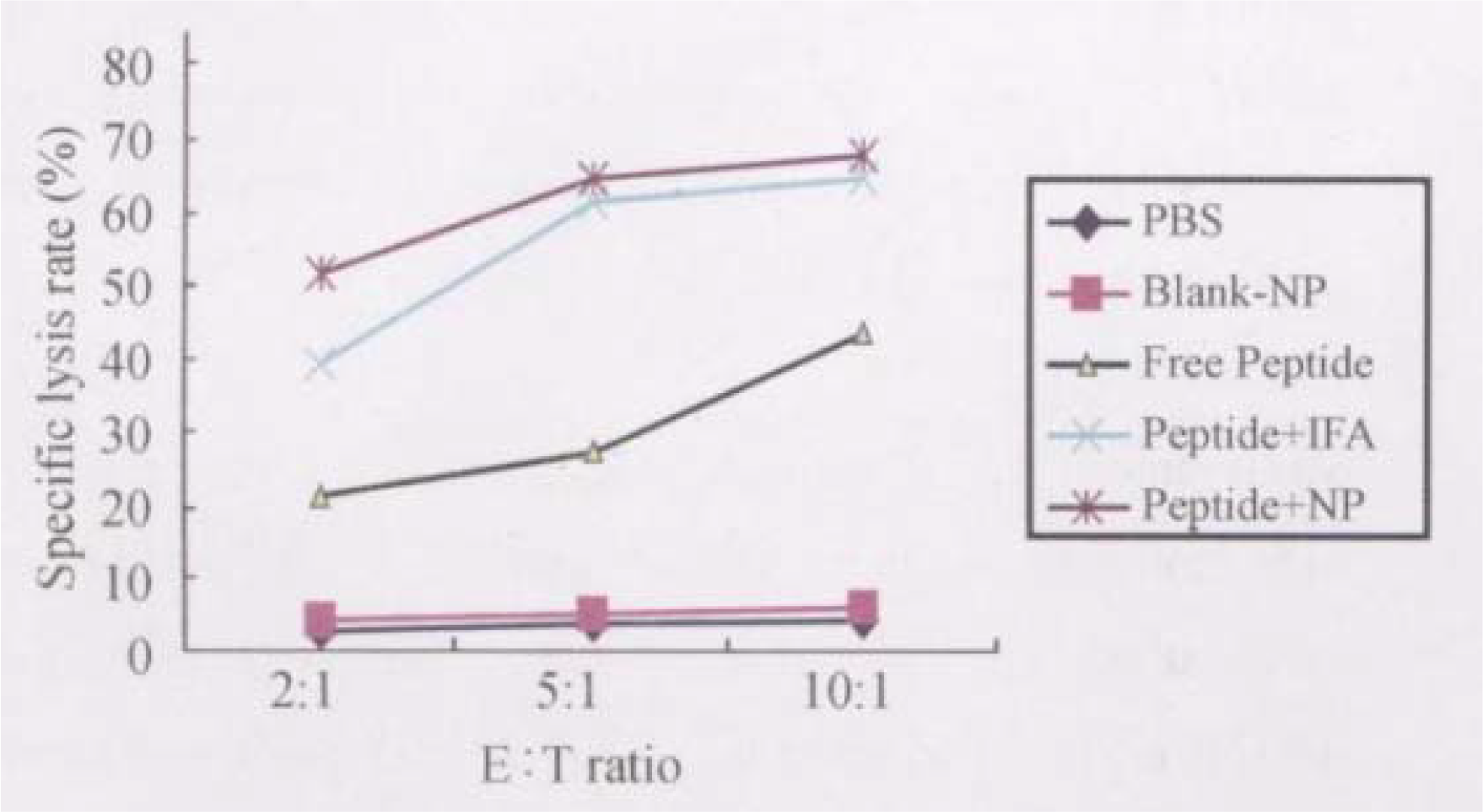
Cytotoxicity study of different treatment groups.

Tumor volume differences between the groups before the group should not statistically significant (P > 0.105).Tumor volume size of each time period in different groups as shown in Figure 6.The tumor inhibition rate was 37.181% in the PBS group, and the difference was statistically significant (P = 0.10125) compared with that in the negative control group (PBS group). 219 survival time compare survival time of mice survival analysis, total median survival of 27 d, the survival time difference in each group no statistical significance (log2rankχ2 =5.1683, P >0105), suggesting that inhibition of tumor growth did not translate into prolong the survival of the limited survival of the effect, See Figure 7.

**Figure 6.**
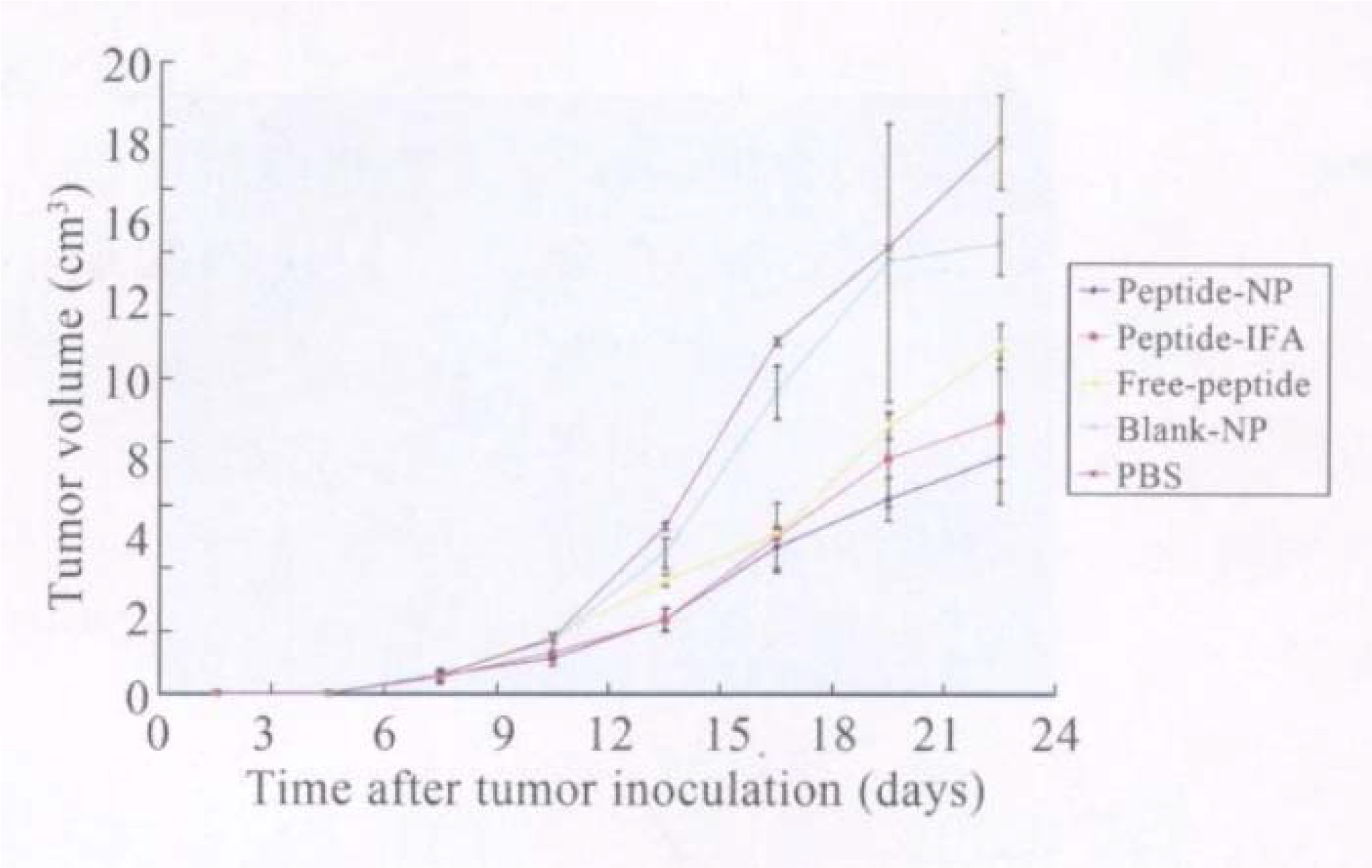
Tumor volume measurement

**Figure 7.**
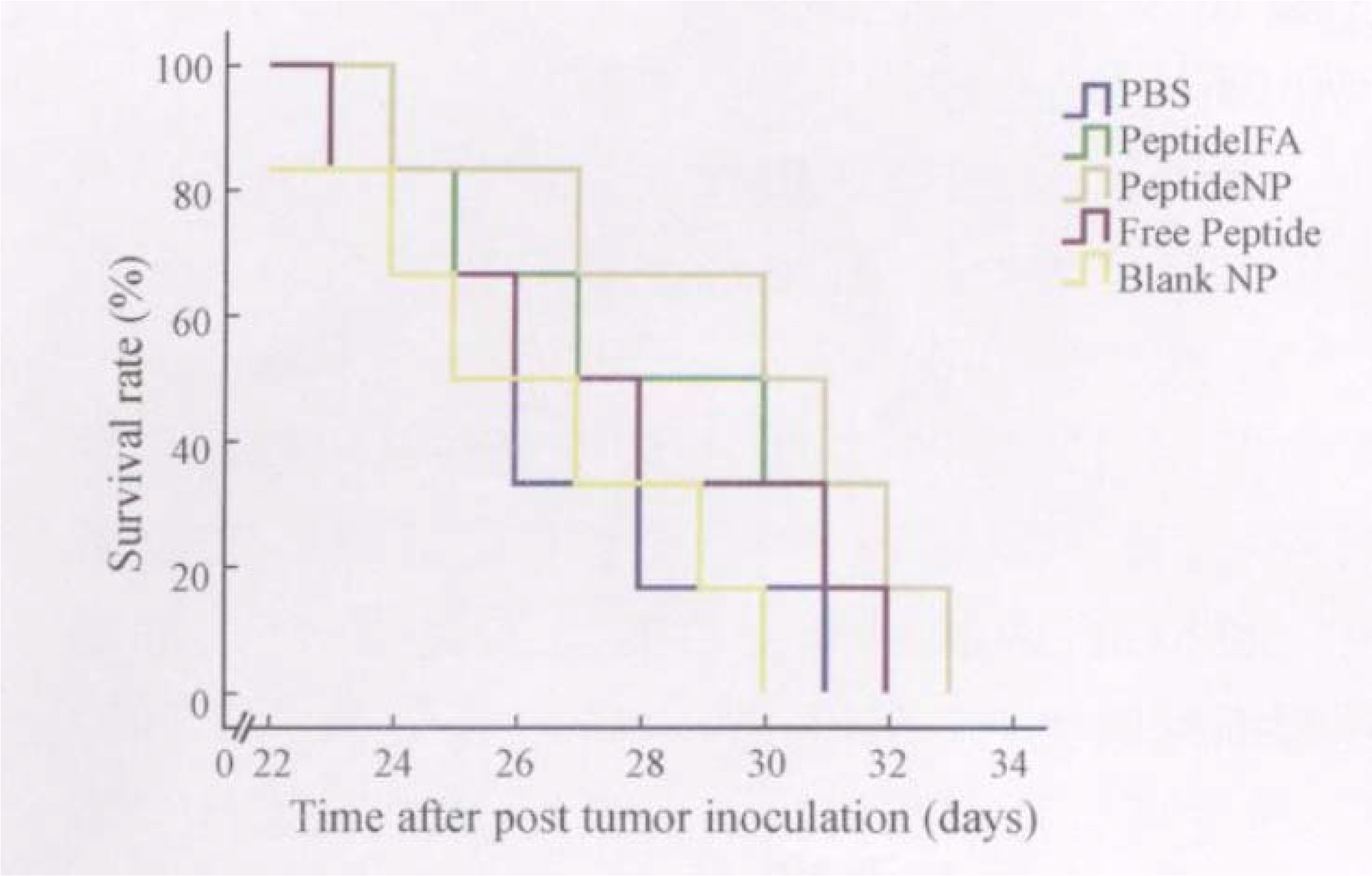
Survival study.

## Discussion

Tumor antigen peptide vaccine discovery, biological response modifiers proposed application and adoptive cell therapy, gene therapy to determine the three-dimensional monument is called tumor biological treatment. Compared with traditional vaccines, tumor polypeptide vaccines have good safety, easy preparation design, easy synthesis of a large number of advantages, which can be used in combination with adjuvants, may induce a strong CD8 + T cell response.In recent years, along with nanotechnology (Yang Jun, etc. 397 1994-2010 China Academic Journal Electronic Publishing House). All rights reserved. (1) the protective effect of the antigen, which can prevent premature decomposition in vivo metabolic process, prolong the antigen residence time in the body, which will help improve the immune Effect; (2) the nanocarrier itself acts as an immune adjuvant, which can assist, expand the non-specific immune response, improve the immune response of the immune system,; (3) nano vaccine particle size is small, with a unique " lymphatic targeting." Traditional polypeptide vaccine mostly single CTL epitope that is mhc2 molecules restricted epitope polypeptide-based [3]. Zwaveling, etc. the study found that the joint th epitope that is mhc2ⅱ molecular-restricted epitope polypeptides can enhance the immune effect.This may be related to the secondary immunological effects of Th cells, the occurrence of double-epitope can inhibit immune tolerance and prolong the duration of antigen presentation. MAGE-3 is a widely expressed antigen in many tumors. Phase I / II clinical trials have demonstrated that MAGE-3 combined adjuvant stimulates specific humoral and cellular immunity, which so far the largest of clinical test (MAGR IT) is also in progress[8, 12-19]. In recent years, the protein carrier material based on amphiphilic graft copolymer is widely studied. Akagi et al. in the experiment using y2pga and L2PAE by self-assembly method to construct amphiphilic nanocarriers found that its biocompatibility, biological safety and biological stability are good, as a protein carrier has certain advantages. Literature also reported that the polypeptide nanometer vaccine can enhance the immune effect, inhibit the growth of tumor cells. Park et al. chitosan was used as the basic raw material, by self-assembly method to build Nano micelles, which can better package polypeptide vaccine. In view of this, chitosan and deoxycholic acid were used as raw materials in this study[14, 20-23]. on the basis of the preparation of nanometer micelles, nanometer micelle units were formed by self-assembly of electrostatic attraction between polypeptide and nanometer micelles, van der Waals force and so on. Through in vitro immunological activity experiments and animal experiments, it was found that the polypeptide vaccine unit with nanomaterial as the loading unit can arouse more effective immune effect than the single pure polypeptide, and compared with the immunoadjuvant group, it has certain advantages. Since the traditional immune adjuvant has certain toxic side effects, the vaccine based on nanomaterials is superior to the traditional immune adjuvant adjuvant immunization method in safety. However, in animal experiments, the vaccine combined with adjuvant group to extend the survival time of mice had no significant advantage, which may be related to the selection of peptide and adjuvant, vaccine dose and immune methods and other factors. Whether it can be combined humoral cellular immune dual way, adjust the vaccine dose and frequency of stimulation interval ways to improve the immune efficacy is one of our future research direction[24, 25]. Nanomaterials as the carrier polypeptide vaccine has broad application prospects, we need to continue to study.

## Conclusion

In this study, we demonstrate that MAGE-3 polypeptide / Chit2 DC nanometer vaccine can effectively activate the immune system and induce anti-tumor immunity. More importantly, multiple doses of micelle vaccines protected mice from tumor challenging. Therefore, future efforts should focus on translating this promising vaccine platform to the clinical settings.

